# Population regulation in semelparous deep-sea squid is driven by ecological conditions in surface waters and whale predation at depth

**DOI:** 10.1101/2024.11.21.624646

**Authors:** Mark Rademaker, Hanna ten Brink, Henk-Jan Hoving, Fleur Visser, Anieke van Leeuwen

## Abstract

Predator-prey interactions present a powerful framework for understanding population regulation in natural systems, including the vast and understudied pelagic deep sea. Oceanic squid are among the most abundant and important species groups in this habitat and function as primary prey for the largest marine top predators, deep-diving cetaceans. At the same time, these cryptic animals are highly data deficient. Insight into the dynamics of deep-sea squid populations and the impacts of predation by top-predators is crucial for designing conservation strategies and developing a general theory of deep-sea ecology. Here, we offer fundamental new insights into how individual life history and predation interact to shape population regulation in deep-sea squid. Using empirical data, we develop a size-structured population model for a highly abundant histioteuthid deep-sea squid. We show that population regulation is driven primarily by conditions experienced as paralarvae in the upper water column, where they face intense resource competition. Relaxation of this competition following ontogenetic migration as juveniles drives exponential-like growth curves. Population dynamics exhibit single-cohort cycles, producing regular ‘seasonal’ patterns in reproduction, even in the absence of any environmental seasonality. Furthermore, we demonstrate that predation by deep-diving cetaceans at different depths can lead to emergent facilitation between top predators. This reveals the complex interdependencies in trophic networks connecting the deep sea with surface waters. Our findings provide critical insights into the ecological functioning of the pelagic deep-sea and their link with surface waters. These insights are urgently needed to better understand and conserve this vast, data-deficient habitat, which endangered deep-diving predators depend on but is under major stress from anthropogenic activities.

## Background

Predator-prey interactions are fundamental to ecological theory [1, 2], and play a critical role in the regulation of populations across marine and terrestrial ecosystems [3, 4, 5, 6]. Predation shapes the dynamics of prey populations through complex feedback mechanisms that depend on prey life histories [7]. Selective predation of specific prey size-classes can alleviate intraspecific competition for resources in non-targeted size-classes, leading to increased individual growth and reproduction, and potentially higher prey biomass [7, 8]. Such emergent, compensatory responses have been empirically shown in systems as diverse as plants [9, 10, 11], fish [12, 13, 8, 14], crustaceans [15], zooplankton [16, 17], and insects [18, 19, 20]. When multiple predator species are present, size-selective predation can even lead to emergent facilitation [21], where one predator species benefits from increased prey availability due to the size-selective predation imposed on the shared prey by another predator [22]. These advances in ecological theory, based on a wide range of systems, can be a powerful tool to gain insights into the processes regulating populations in critical yet understudied ecosystems.

The pelagic deep-sea is the largest and least studied natural habitat on earth and represents an important reservoir of species diversity and biomass [23, 24]. Deep-sea squid form a key link in pelagic deep-sea food webs by foraging on plankton and (micro)nekton and functioning as the primary prey for a wide range of marine top predators [25, 26, 27]. The largest of these predators are toothed whales, a group of cetaceans specialized in diving to extreme depths [28, 29]. The predation of deep-sea squid by deep-diving cetaceans exemplifies the direct trophic links that exist between the pelagic deep sea and surface waters. Many deep-diving cetacean species co-occur and forage on similar prey communities of deep-sea squid [30, 31]. Recent studies indicate that deep-diving cetaceans limit competition for prey by targeting distinct depth ranges [31]. Since commonly predated deep-sea squid are distributed by size across different depths [32, 33], this suggests that deep-diving cetaceans impose size-selective predation mortality on deep-sea squid. How prey populations respond to such size-selective mortality is dependent on their life history [7]. To understand the consequences of predation by deep-diving cetaceans, we must therefore first better understand the life history and internal population regulation of deep-sea squid.

The deep-sea squid genus *Histioteuthis* is among the most abundant and relatively well-studied groups of deep-sea squid [34, 35]. They are also among the most reported squid prey of deep-diving cetaceans [30, 31, 36]. The few studies available suggest that *Histioteuthis* exhibit relatively fast growth rates, semelparous reproduction, and a short lifespan of 1-2 years [34, 32]. Like other deep-sea cephalopods, growth in *H*. reversa follows an exponential pattern without an upper plateau [37, 38]. The mechanism underlying this unusual pattern of faster growth with increasing size is unknown, but has been reported repeatedly for squids. It might occur because across deep-sea cephalopods the metabolic maintenance costs (per gram body mass) decrease with increasing inhabited depth [39, 40]. In species with ontogenetic vertical migration, bigger individuals inhabiting larger depths might therefore have relatively more energy available to allocate towards growth. Alternatively, the exponential growth patterns might reflect the under sampling of the largest (mature) size classes in surveys [32, 38]. There are indications that maturation in *Histioteuthis* follows seasonal patterns [32, 34]. These seasonal patterns could arise from individuals postponing the ripening of their gonads to synchronously reproduce when conditions are optimal; a phenomenon observed in several fish and squid species [41, 42, 43]. Alternatively, maturation and reproduction might occur year-round, but the seasonal patterns could result from a single cohort of offspring, born under optimal conditions, outcompeting earlier cohorts. Such single cohort cycles have been described in fish [44, 45] and krill [46]. Single cohort cycles leave a strong imprint on population dynamics by causing periodic shifts in the dominance of juvenile and mature adult individuals. Enhancing our mechanistic understanding of growth and maturation in *Histioteuthis* will therefore provide fundamental insight into how their populations are regulated, which is of direct relevance for conserving their deep-diving cetacean predators.

Here we use empirical data to develop a size-structured population model of the hyperabundant and trophically important deep-sea cephalopod, the reverse jewel squid (*H. reversa*). We use this mathematical model to mechanistically understand the processes underlying observed patterns of individual growth and maturation in deep-sea squid, and the population level dynamics emerging from these individual level processes. Such understanding then ables testing of the hypothesis that depth-specific predation by deep-diving cetaceans results in 1. deep-sea squid biomass overcompensation and 2. emergent facilitation between co-existing predators. We use the deep-diving cetaceans Risso’s dolphin (*Grampus griseus*) and goose beaked whale (*Ziphius cavirostris*) as model predators. These species both forage on *H. reversa*, and while they co-occur, they forage at distinct depth ranges [31]. The outcomes of this study will provide fundamental new insights into how deep-sea squid populations are regulated and affected by predation from higher trophic levels. This presents a key step to forming a more general theory of the functioning of pelagic deep-sea ecosystems across trophic levels. Next to its fundamental importance, there is an urgent need for such a theory to be able to better predict and mitigate the current strong impacts of human-induced changes in this immense but understudied ecosystem.

## Methods

### Model summary

We model the *H. reversa* population using a size-structured population model that describes dynamics based on the individual level processes of growth, reproduction, and mortality [47]. Model parameters and references can be found in Table 1 and a conceptual representation of the model is presented in Figure 1. In our model system individual *H. reversa* develop through three life stages: paralarvae, juveniles, and (mature) reproducing adults. The egg-stage of *H. reversa* is not modeled, but egg mortality and energy conversion efficiency from egg to hatchling paralarvae stage are accounted for. Paralarvae forage on zooplankton resources in the upper water column and switch to forage on a mix of nekton and crustaceans in the deep-scattering layer when reaching juvenile size. As juveniles increase in size they move to deeper habitats and gradually transition to foraging on a second nekton resource in deeper layers (Figure 1). The spatial separation between smaller juveniles in the deep-scattering layer and maturing individuals and adults that remain in the deepest waters throughout the diurnal cycle is implicitly accounted for through separation of resource types. During development, juveniles allocate an increasing fraction of their ingested energy to the production of gonads. Once the gonads of an individual have reached a threshold level, the individual matures into the reproducing adult stage and immediately releases all its eggs. At this stage we assume individuals stop feeding and only reproduce, eventually dying of starvation post-reproduction. Below, we provide a mathematical definition of the life history functions in our model and the underlying biological assumptions in relation to individual growth, mortality, and reproduction.

**Table 1.** Parameters and default values for the *H. reversa* model. Calculations to convert the values and units as presented in the source references to those listed in the table are presented in the online data repository.

| Parameter | Default value | Unit | Description | Source species & Ref. |
| --- | --- | --- | --- | --- |
| $\delta_i$ | 0.1 | D | Resource turnover rate | Modelling assumption [51] |
| $R_{1\max}, R_{2\max}, R_{3\max}$ | Variable | $\text{g}\cdot\text{L}^{-1}$ | Maximum biomass density of resource 1,2, 3 | - |
| $\sigma$ | 0.42 | - | Conversion efficiency | *I. Illecebrosus* [96] |
| $a$ | 0.6 | $\text{L}^{-1}\cdot\text{g}^{-1}\cdot\text{d}^{-1}$ | Maximum mass-specific attack rate | Modelling assumption [97] |
| $h$ | 1 | $\text{day}^{-1}$ | Mass-specific handling-time | Modelling assumption [97] |
| $\tau$ | 0.1 | $\text{day}^{-1}$ | Mass-specific maintenance rate (depth dependent) | *H. hoylei*, *H. heteropsis* [39, 98, 99] |
| $\mu_b$ | 0.028 | $\text{day}^{-1}$ | Background mortality rate paralarvae | *H. reversa*, *H. bonelli* [37, 100] |
| $\mu_j$ | 0.016 | $\text{day}^{-1}$ | Background mortality rate juveniles & adults | *H. reversa*, *H. bonelli* [37, 100] |
| $\mu_c$ | Variable | $\text{day}^{-1}$ | Additional predation mortality by *Z. cavirostris* | - |
| $\mu_g$ | Variable | $\text{day}^{-1}$ | Additional predation mortality by *G. griseus* | - |
| $\theta$ | 0.5 | - | Egg to offspring energy conversion efficiency | Modelling assumption [51] |
| $\beta_{\min}$ | 76.8 | g | Minimum reproductive buffer biomass | *H. reversa* [101] |
| $S_b$ | 0.0016 | g | Paralarvae size at birth | *H. reversa* [32, 101] |
| $S_j$ | 2.1 | g | Paralarval to juvenile transition size | *H. reversa* [34, 102] |
| $S_{\max}$ | 811 | g | Maximum size | *H. reversa* [32, 34] |
| $S_m$ | 93 | g | Maturation size | *H. reversa* [32, 102] |
| $S_{\text{shallow}}$ | 2.1 | g | Size at start depth migration | *H. reversa* [34, 102] |
| $S_{\text{deep}}$ | 405 | g | Size at half the maximum depth migration | *H. reversa* [32] |

**Figure 1.**
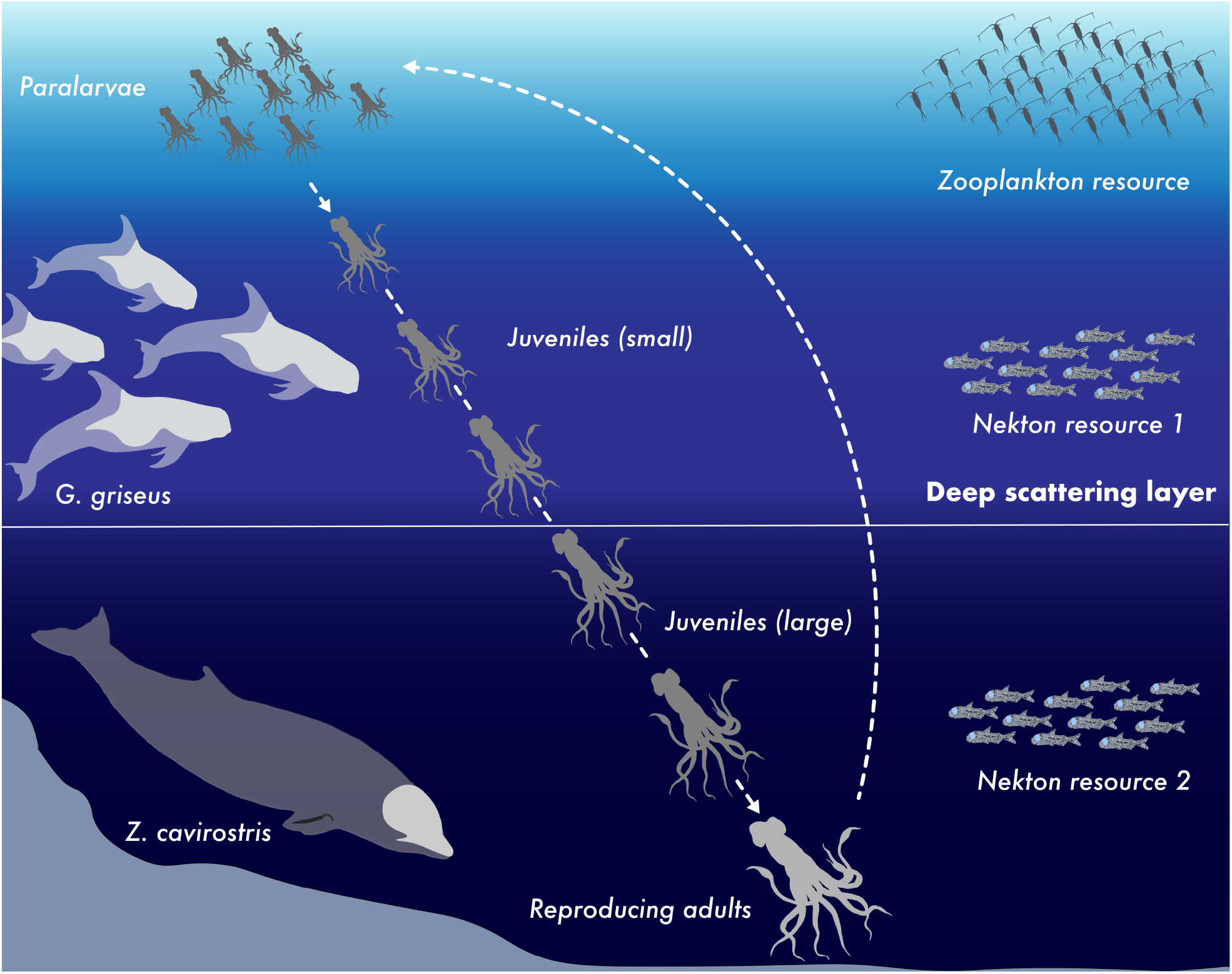
Conceptual representation of deep-sea squid life history, predators, and prey resources. The model study system with prey (squid) and predators (two deep-diving cetacean species, *G. griseus* and *Z. cavirostris*). Squid paralarvae remain in the upper water layer where they forage on zooplankton. Juvenile squid undergo ontogenetic migration by which they inhabit increasingly large depths with increasing size and are subjected to size-selective predation by two deep-diving cetacean predators at different depth ranges. Straight dashed arrows indicate squid progression to subsequent life-stages through growth, while the curved dashed arrow indicates biomass flow through reproduction. The horizontal white line represents the implicit separation in the model between smaller-sized juveniles and their prey in the deep-scattering layer, and maturing juveniles, adults, and their prey, that remain in deeper layers.

### Life history functions and assumptions

We model three squid life stages that are distinguished based on a threshold body size. Individuals post-hatching belong to the paralarvae stage. Paralarvae feed upon a zooplankton resource with density *R*_1_. Paralarvae switch to the juvenile stage at body mass *S*_j_. Juveniles initially forage on nekton resource *R*_2_, and gradually shift to foraging on nekton resource *R*_3_ as they grow larger (representing the use of deeper ocean habitats with ontogeny). In addition to somatic growth, juveniles build up reproductive biomass. After reaching a reproductive buffer mass of β_min_, we classify individuals as adults. The build-up of this reproductive buffer depends on the body size of the individual and the available food. We assume that adults immediately release their reproductive buffer after reaching the reproduction threshold. Due to the cessation of feeding following reproduction, adults die of starvation. Several of the model’s life history functions depend on the relative depth, *ρ*, inhabited by an individual. A *ρ* value of 0 indicates that an individual remains at the surface and a value of 1 indicates that individuals have reached the maximum depth (Supplementary Figure 1). This relative depth depends on individual body size *S*, and follows a continuous, piecewise-differentiable, sigmoid function that ranges between 0 and 1 [48], such that

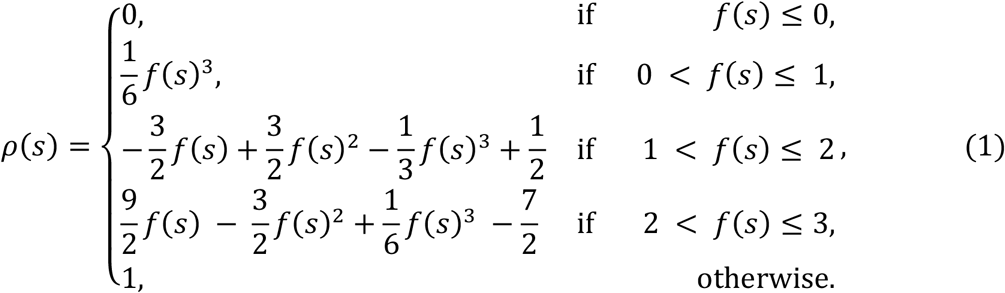

Individuals start migrating downwards at a body size of *s* = *S*_Shallow_, and reach half the maximum depth at a body size of *s* = *S*_deep_. In equation 1, the function *f*(*s*) equals

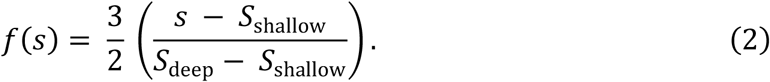

### Food intake

Individuals forage on multiple resources with densities *R*_*i*_ (with *i* = 1, 2, 3) following a holling type II functional response. Paralarvae, with body mass *S* < *S*_j_, feed upon zooplankton (*R*_1_) only, while juveniles can feed upon two different types of nekton resources (*R*_2_ and *R*_3_) depending on their inhabited depth *ρ*(*S*). We assume that reproducing adults do not feed. Mass-specific food intake of paralarvae and juveniles, *I*(*S, R*_1_, *R*_2_, *R*_3_), equals

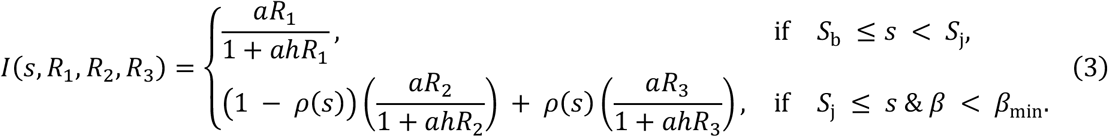

In this equation, parameter *a* represents the mass-specific attack rate and parameter *h* the handling time.

### Biomass production

Ingested food is assimilated with efficiency *σ*. Individuals use assimilated energy first to cover their mass-specific maintenance costs *τ*. We assume that these maintenance costs decrease linearly with the inhabited relative depth, *ρ*(*S*). The mass-specific net-biomass production rate of an individual, *ν*(*s, R*_1_, *R*_2_, *R*_3_), is then given by the difference between energy intake and maintenance costs following

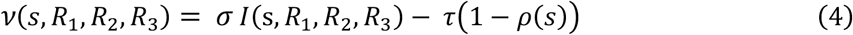

We also examine the scenario in which there is no decrease in mass-specific maintenance costs with increasing inhabited relative depth, in which equation 4 reverts to *σ I*(*s, R*_1_, *R*_2_, *R*_3_) ™ τ. When energy uptake is insufficient to cover maintenance costs (*σ I*(*S, R*_1_, *R*_2_, *R*_3_) < *τ*(1 − *ρ*(*s*)), individuals experience starvation mortality, *μ*_*S*_, equal to -*ν*(*s, R*_1_, *R*_2_, *R*_3_) on top of their background and predation mortality (see section on mortality below). Note that due to our assumption of non-feeding adults, adults always starve with a per capita rate of *τ*(1 − *ρ*(*s*))*s*, which equates −*ν*(*s, R*_1_, *R*_2_, *R*_3_) in this case.

As individuals transition from the paralarval to the juvenile stage and grow towards maturation, they invest a decreasing proportion, *κ*(*s*), of net biomass production into somatic growth, and an increasing proportion, (1 - *κ*(*s*)), into their reproductive buffer. The fraction *κ*(*s*) is modeled as a smoothly decreasing function of body size following

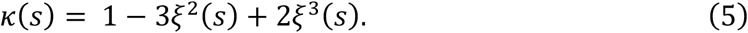

In this equation, *ξ*(*s*) equals

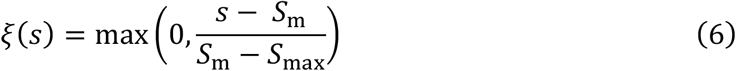

where parameter *S*_m_ equals the minimum maturation body size and parameter *S*_max_ the theoretical maximum body size individuals can reach.

### Fecundity

When the reproductive buffer *β* of an individual reaches the level *β*_min_, the reproductive storage of an individual is immediately transformed into newborn paralarvae. The number of paralarvae an adult produces is given by

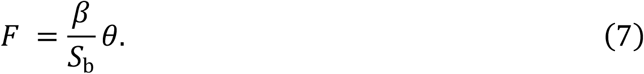

Where *θ* indicates a conversion of efficiency to account for mortality during the egg stage due to predation and disease.

### Mortality

Background mortality rates for *H. reversa* are life-stage specific, with a high constant background mortality for paralarvae in the upper water column, and a lower mortality rate for juveniles. In addition to background mortality, individuals may experience the above-described starvation mortality, *μ*_*S*_(*S, R*_1_, *R*_2_, *R*_3_), and predation mortality imposed by deep-diving cetaceans. Total mortality rate is given by

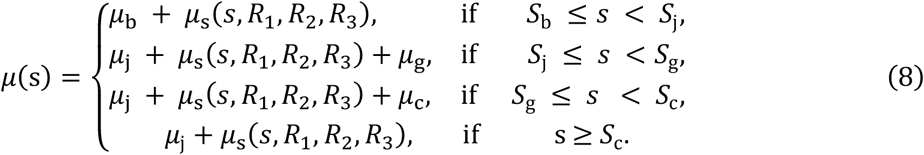

In this equation, *μ*_g_ equals the additional predation mortality imposed by *G. griseus* and *μ*_c_ the additional mortality imposed by *Z. cavirostris*. We assume that *G. griseus* feeds on individuals at shallower depths, characterized by a body mass between *S*_j_ and *S*_g_, and that *Z. cavirostris* feeds on individuals at larger depths, characterized by a body mass between *S*_g_ and *S*_c_. Hence, the depth-dependent predation by the two predators can be represented by their specialization on two non-overlapping size ranges of *H. reversa*. We believe this model assumption to be justifiable in the context of our analysis because of (1) the strong relationship observed between size and inhabited depth of *H. reversa* across the diurnal cycle [32], and (2) the clear separation in main foraging depth observed between *G. griseus* and *Z. cavirostris*, respectively at 200-600 m and 950 m to the bathyal sea floor [31]. We acknowledge that there could always be some level of context-dependent overlap in prey selection between both predators. For example, due to spatial or temporal variation in prey availability at different depths, or the structure of the predator population (size of the pod, age structure, and body condition). However, these dimensions of variation fall outside the scope of the general mechanisms we aim to investigate in this study.

### Population dynamics

Since we assume in our model that adults release all their eggs at once, all offspring produced at a reproductive event are grouped into a single cohort and assumed to experience identical environmental conditions and hence to develop at the same rate. The dynamics of each cohort *i* ∈ N can be followed numerically by integrating a set of three Ordinary Differential Equations (ODEs). These ODEs keep track of the number of individuals in the cohort, *c*_*i*_, their body mass, *S*_*i*_, and their reproductive buffer, *β*_*i*_. When a cohort of adults reproduces, a new cohort of paralarvae is added to the population, which results in three additional differential equations describing the population dynamics. Cohorts pass through three life stages: the paralarval (L), juvenile (J), and adult stage (A). Transitioning between life stages depends on cohort size (*S*_*i*_) and cohort buffer (*β*_*i*_). The dynamics of the density, body mass, and reproductive buffer in paralarvae and juveniles depend on the amount of food they encounter and can be described by the following set of ODEs:

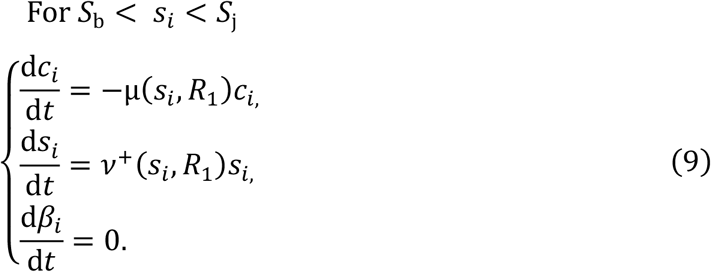

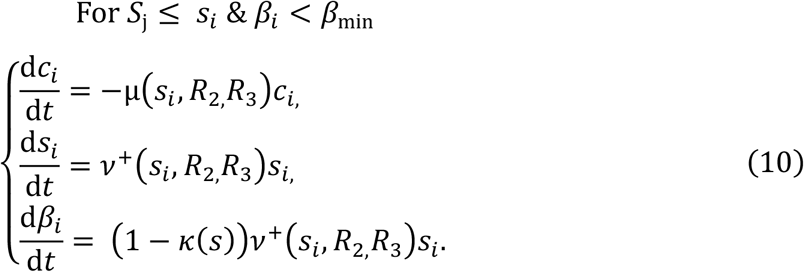

In these equations we use the notation *ν*^+^(*s*_*i*_, *R*_1,_*R*_2,_*R*_3_) to indicate the mass-specific net-biomass production rate restricted to positive values only. Whenever the reproductive buffer of the oldest juvenile cohort with index *i = m* reaches the threshold value *β*_m_ = *β*_min_, at time *t* = *t*_repro_, a reproduction event occurs. This juvenile cohort becomes an adult cohort, equal in number and size to the cohort just before the reproductive event 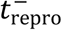. These adults reproduce immediately and a new cohort with index 0 is formed from the biomass stored until just before reproduction 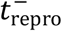. The buffer of the reproducing cohort is set to 0 and all cohorts (both reproducing and non-reproducing) are renumbered. The changes are described by the following sets of equations:

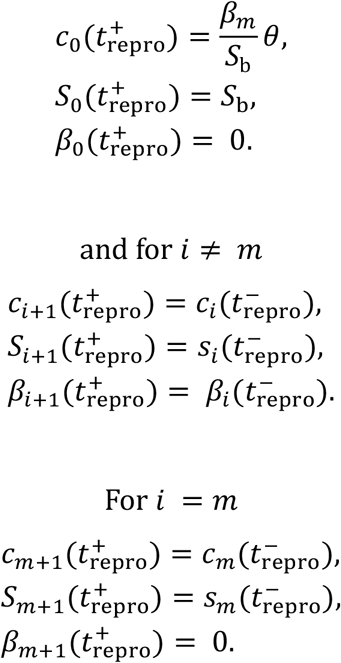

The spent adults no longer feed and their dynamics are thus no longer dependent on food densities. Adults only starve and their dynamics can be described by the following set of ODEs:

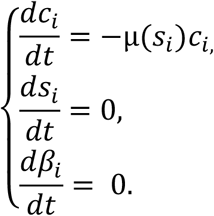

### Resource dynamics

Our model includes three unstructured resources expressed in grams per unit volume. The biomass dynamics of all three resources are described by a turnover rate δ_*i*_, maximum resource density *R*_*i*,max_, and consumption by *H. reversa* life stages (eqn. 19-21). The zooplankton resource dynamics can be expressed as

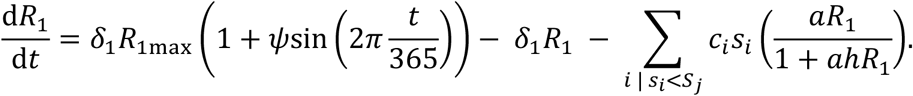

The summation term in this equation relates to the resource ingestion of all paralarval cohorts. The productivity of the zooplankton resource is assumed to follow the seasonality of primary production in surface waters and is modeled through a sinusoidal function with a period of 365 days and amplitude *ψ* ranging from 0 < *ψ* < 1. Ecologically, a *ψ* value of 0 means there is no seasonality and resource productivity is spread evenly over the year. In contrast, *ψ* values ranging between 0 < *ψ* < 1 indicate a range of weak to strong seasonality where peak resources productivity occurs in an increasingly narrow time window within the year. The total resource production within a year is the same irrespective of the value of *ψ*. We assume the nekton resources *R*_2_ and *R*_3_ in the deep-sea do not follow the seasonality of primary productivity in surface waters like *R*_1._ Their dynamics can therefore be expressed as

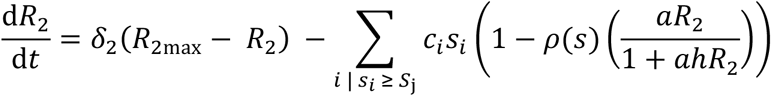

and

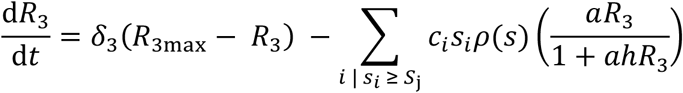

The summation term models the ingestion of the resource over all juvenile cohorts.

### Model parameterization and analysis

We parametrized the life history functions for *H. reversa* based on scientific literature (Table 1). We were able to derive most life history parameters directly from published empirical studies on *H. reversa* (References in Table 1). When this information was not available, we used studies of closely related species (*H. hoylei, H. heteropsis, H. bonelli*), where possible. The life history functions and the set of ODEs describing the dynamics of *H. reversa* life stages were coded in C programming language and numerically integrated using the escalator boxcar train (EBT-train) method [49], using the EBT-tool [50]. We ran time-series of *H. reversa* population dynamics over 1,000,000 time-steps, where each individual time-step represents one day. Each model run was initialized with 6 differently sized cohorts, each containing 100 individuals. After ensuring that the population dynamics had reached a stable attractor, we extracted the final 3,000 time-steps from the model output to visualize population level abundance, biomass dynamics, and individual (cohort) growth. We ran the model with and without depth-dependent metabolic costs to examine its effect on population dynamics and individual growth. Next, we examined how sensitive population and individual level dynamics were to changes in life history and resource parameters, by means of bifurcation analysis. To this end we ran an iterative time-series analysis with intervals of 100,000 time-steps. At the start of each new iteration, we systematically increased or decreased the value of a focal bifurcation parameter with a marginal amount (plus or minus 0.001 as step-size). We studied the effect of changes in resource productivity (*R*_1max_, *R*_2max_, *R*_3max_), background mortality (*μ*_b_, *µ*_j_), depth migration (*S*_deep_), resource seasonality (*ψ*), and size-selective predation mortality (*μ*_g_, *µ*_c_), in the bifurcation analysis.

## Results

### Life-stage specific individual growth and maturation

Individual squid showed an exponential-like growth curve (Fig. 2a). However, rather than a constant mass-specific growth rate as expected under true exponential growth, we found an underlying pattern of a sharp decrease in mass-specific growth early in life (for paralarvae), followed by an increase in mass-specific growth with increasing size in juveniles (Fig. 2b). This pattern of exponential-like growth is not driven by reductions in mass-specific maintenance costs with increasing depth (Supplementary Figure 2). Rather, it can be traced to the high resource competition experienced in the paralarval life stage, limiting energy intake, and the subsequent relaxation of resource competition and availability of multiple resources following ontogenetic vertical migration in the juvenile stage (Fig. 3; these findings are discussed in detail in the next section). Finally, we find the pattern of exponential-like growth curves and increased mass-specific growth with juvenile size to be robust with respect to changing model parameters. Growth-patterns only change under very extreme and limited ecological conditions (Supplementary Figure 3.1). Specifically, a qualitative reversion, where mass-specific growth decreases with increasing size, occurs only when high paralarval mortality is coupled with both low productivity of the secondary nekton resource in deeper layers and an earlier switch to this resource (at 200 g body mass instead of the default 400 g)

**Fig. 2.**
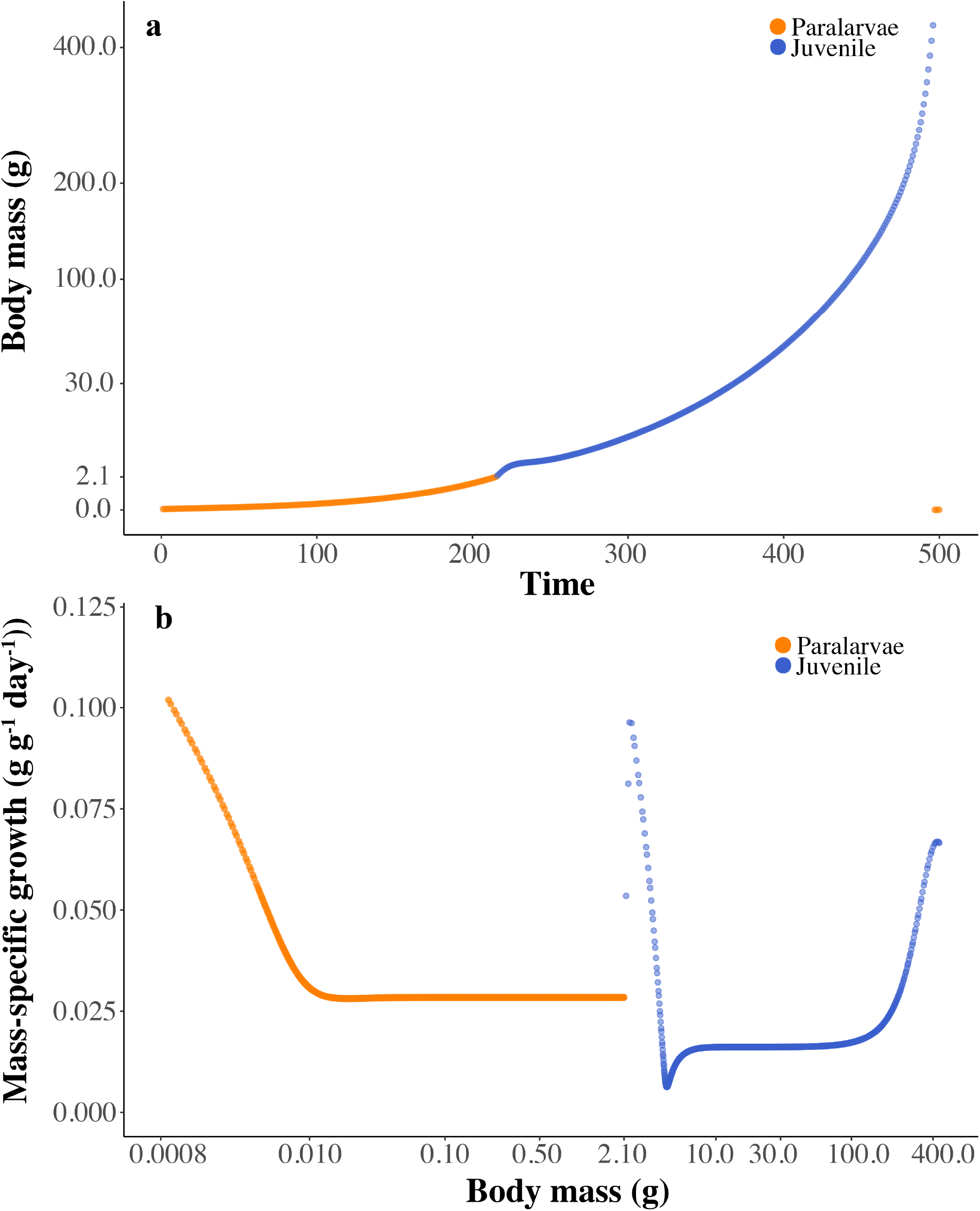
Growth and maturation of deep-sea squids. Individual growth curve (a) and mass-specific growth (b) for default parameters (Table 1). As growth spans seven orders of magnitude from egg to maturation we applied a square root scaling to the y-axis of (a) to visualize growth patterns across the life cycle without linearizing them (as under the log transformation). An untransformed version of (a) is provided in Supplementary Figure 3.2; showing the square-root scaling did not affect the qualitative growth pattern (exponential).

### Population dynamics follow cohort cycles

The modelled population dynamics of *H. reversa* are characterized by single cohort cycles (Fig. 3). These cycles start with a high reproductive inflow into the paralarval life stage (Fig. 3a). This influx leads to intense (intra-cohort) resource competition among the newborn paralarvae, reflected by a sharp drop in zooplankton resources at the birth of each new cohort (Fig. 3c). This drop in resources causes high starvation mortality and initial steep declines in paralarvae densities (Fig. 3a). As densities of paralarvae decline, starvation mortality decreases due to relaxation of competition. However, competition for zooplankton between surviving paralarvae is still sufficiently high to limit their growth, resulting in the accumulation of biomass in the late paralarval stage (near horizontal part of the orange curves in Fig. 3b). This bottleneck is lifted when individuals transition to the juvenile stage and gain access to the first (nekton) resource, leading to a rapid increase in growth and subsequently small juvenile biomass (Fig. 3b). The juveniles soon deplete the first (nekton) resource to just above the critical resource density needed to support their maintenance costs (dotted line, Fig. 3c), leading to the formation of a second biomass bottleneck (near horizontal part of the blue curves in Fig. 3b). This bottleneck is lifted as individuals move to a deeper habitat following ontogenetic vertical migration and gain access to the second (nekton) resource (Fig. 3b). The densities of the remaining large juveniles at this stage are low and these individuals experience little resource competition (Fig. 3a), allowing them to quickly increase in biomass until they reproduce (Fig. 3b). This results in the creation of a new, abundant cohort of paralarvae, which restarts the above described cycle.

**Fig. 3.**
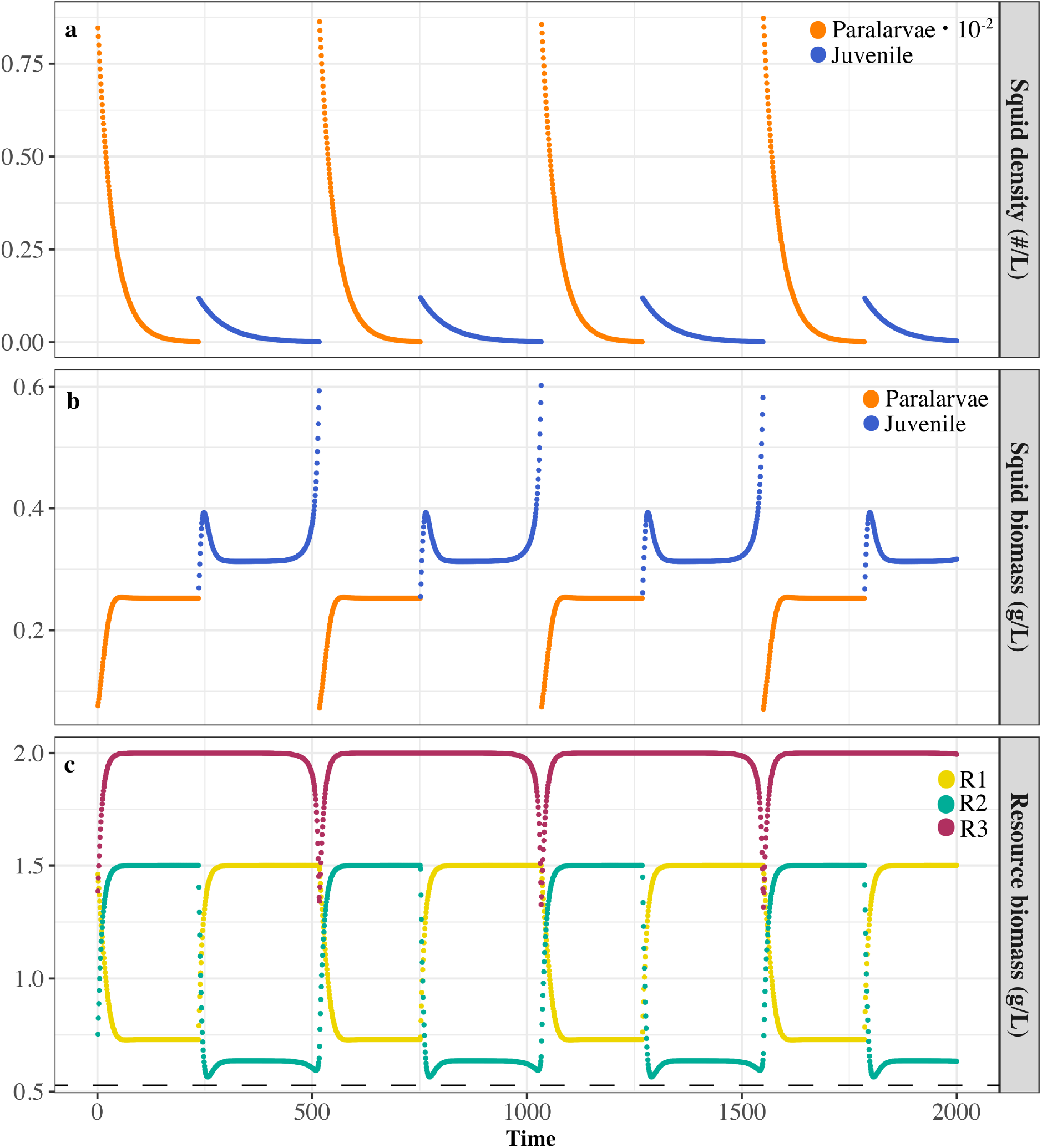
Single-cohort dynamics and paralarval bottlenecks in deep-sea squid populations. Dynamics of squid density (a), squid biomass density (b), and resource biomass density (c) as a function of time. Population dynamics of squid are characterized by single cohort cycles with bottlenecks in the paralarval stage and early juvenile stage due to (intra-cohort) resource competition. Juvenile cohorts (blue) suppress the first nekton resource (green) to just above the critical energy density required to cover metabolic costs (black dotted line). The bottleneck is lifted once juveniles gain access to the secondary nekton resource (red line). Parameters *R*_1max_, *R*_*2*max_, *R*_3max_ were set at 1.5, 1.5, and 2.0 g/L respectively. Other parameters as in Table 1.

We find the population dynamics of *H. reversa* to be generally robust with respect to changes in life history and resource model parameters (Supplementary Figure 4.1-4.6). Over the ranges of these parameters, the regulatory mechanisms remain the same and the population remains characterized by single cohort cycles (e.g. Supplementary Figure 5). However, the total population biomass and reproductive output of the population are sensitive to changes in paralarvae mortality; either imposed directly (Supplementary Figure 4.1), or indirectly through seasonality in zooplankton resources (seasonal effects discussed in the next section).

### Seasonality of zooplankton production affects population dynamics

The population structure of *H. reversa* as seen in the single cohort cycle dynamics can change depending on the seasonality in our model. Strong seasonality in zooplankton productivity (0.5 < *ϕ* < 0.9) reduces total population biomass and switches the squid population biomass from being dominated by juveniles to dominated by paralarvae (Fig. 4a). The underlying mechanism is that high seasonality results in high mortality of paralarvae born outside peak resource conditions. This strongly reduces the competition between surviving paralarvae, illustrated by the fast growth and strong reduction in time spent in the paralarval stage (Fig. 4b). Conversely, the total time spent in the juvenile life stage now increases (Fig. 4b), and the total reproductive output of the population is reduced (Fig. 4c). This indicates that there is relatively more competition between individuals near maturation sizes. Despite these changes in population structure, the population dynamics remain characterized by single-cohort cycles (Supplementary Figure 5).

**Fig. 4.**
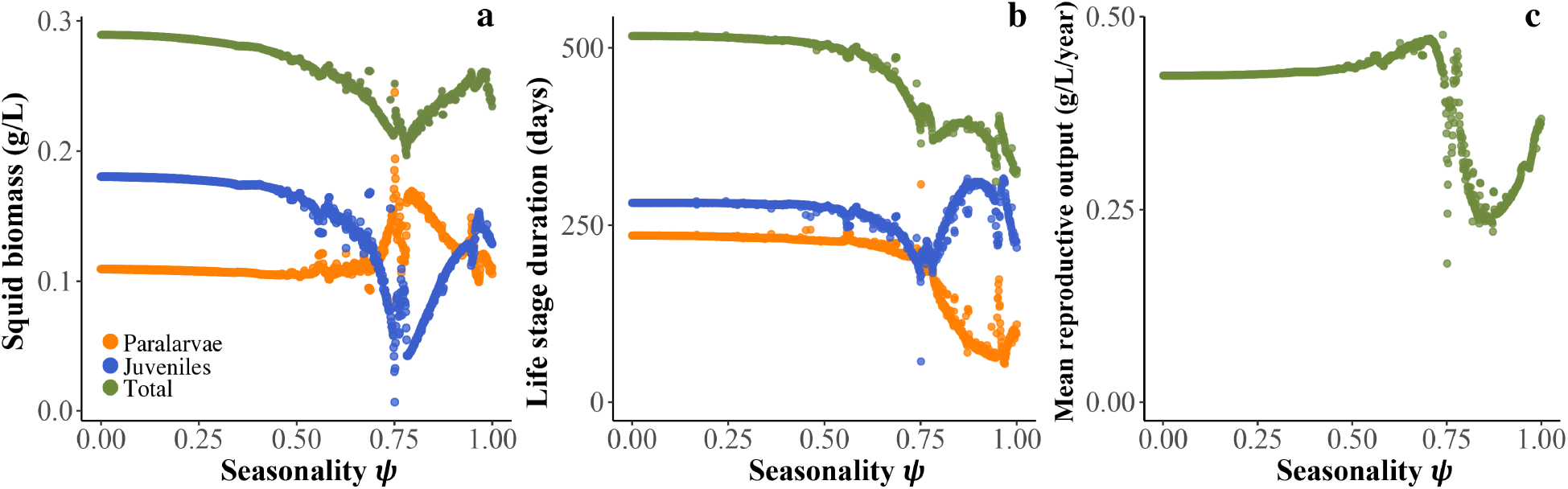
Seasonality drives dominance of life-stages and reproductive output in deep-sea squid *H. reversa*. (a) Average biomass of the three life stages, (b) duration of each life stage, and (c) mean reproductive output rate as a function of seasonality in paralarvae resource productivity. Apart from changing the seasonality, default parameters were used (Table 1). Seasonality 0 = none, 1 = maximum. Parameters *R*_1max_, *R*_*2*max_, *R*_3max_ were set at 1.5, 1.5, and 2.0 g/L respectively. Other parameters as in Table 1.

### Depth-selective predation affects population dynamics and predator diversity

Size-selective predation on *H. reversa* can increase prey availability for other predators. This size-dependent phenomenon has been previously named emergent facilitation [21, 22]. Predation of *Z. cavirostris* on larger juvenile and adult squid at greater depths increases the biomass of smaller, juvenile squid inhabiting the shallower depth range targeted by the cetacean predator *G. griseus*. This positive feedback by *Z. cavirostris* on the prey availability for *G. griseus* persists over a large range of predation pressures (Fig. 5a). The underlying mechanism is that the size-selective predation of large juveniles by *Z. cavirostris* directly suppresses the biomass of these size classes (Fig. 5a light orange line). As a result, fewer large individuals mature and reproduce, resulting in less biomass inflow into the paralarval stage (Supplementary Figure 6.1). The reduced reproduction due to predation means that paralarvae experience less competition for resources, resulting in faster growth and a faster transition to the juvenile stage (Supplementary Figure 6.1). This results in biomass accumulation in the small juvenile size-classes that inhabit shallower depths (Fig. 5a light blue line). As a result of this mechanism, predators like *G. griseus*, which forage on these smaller size-classes, have more prey available.

**Fig. 5.**
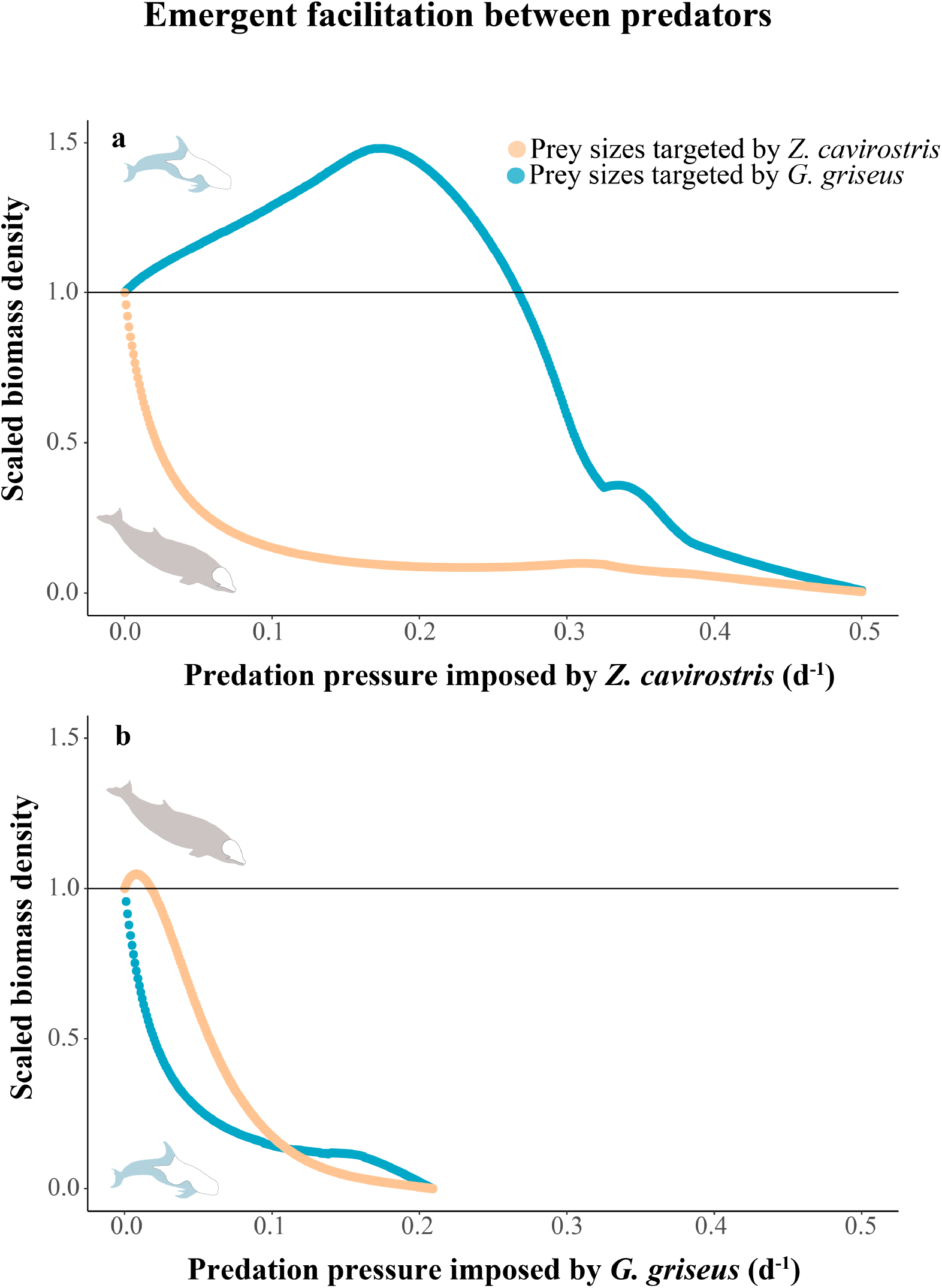
Emergent facilitation between deep-diving cetacean predators imposing size-selective predation on their deep-sea squid prey. The scaled biomass density of squid prey size classes preferred by *Z. cavirostris* (orange-pink line) and *G. griseus* (blue line) are presented as a function of the predation pressure by each predator (a, b). The scaling is relative to the average prey biomass density when there is no predation by either predator (horizontal line at 1.0 on the y-axis). Emergent facilitation occurs when prey biomass density exceeds 1.0 for a predator as a function of increased predation pressure by another predator. Parameters *R*_1max_, *R*_*2*max_, *R*_3max_ were set at 1.5, 1.5, and 2.0 g/L respectively. Other parameters as in Table 1.

The predation of smaller squid at shallower depths by *G. griseus* can slightly increase the biomass of larger squid at greater depth targeted by *Z. cavirostris* (Fig. 5b); but only under low predation pressure. For intermediate and high predation pressure, *G. griseus* negatively affects prey availability for *Z. cavirostris*. When predation pressure of *G. griseus* is low, the reduced competition between remaining juveniles means they can grow faster and larger, increasing the biomass available in the larger size classes targeted by *Z. cavirostris* (Fig. 5b, light blue line). However, this benefit falls away when predation pressure by *G. griseus* increases, as the relative increase in large juvenile prey biomass no longer outweighs reduction in the number of juvenile individuals reaching this stage.

### Increased scope for emergent facilitation in seasonal environments

The overall qualitative pattern of a positive effect of predation by *Z. caviorostris* on prey availability for *G. griseaus* is similar between seasonal and aseasonal environments. Apart for a small range of low predation pressures, we find that the strength of the positive effect of emergent facilitation is approximately twice as strong in seasonal environments (Fig. 6a) compared to aseasonal environments (Fig. 5a). The interaction between seasonality and predation by *Z. cavirostris* amplifies the emergent facilitation effect by increasing the biomass buildup in small-juvenile size classes (Supplemental Figure 7.1 compared to 6.1). The increased biomass of small juvenile size-classes means they experience more competition, grow slower, and remain in the size-range at which they are vulnerable to predation by *G. griseus* longer.

**Fig. 6.**
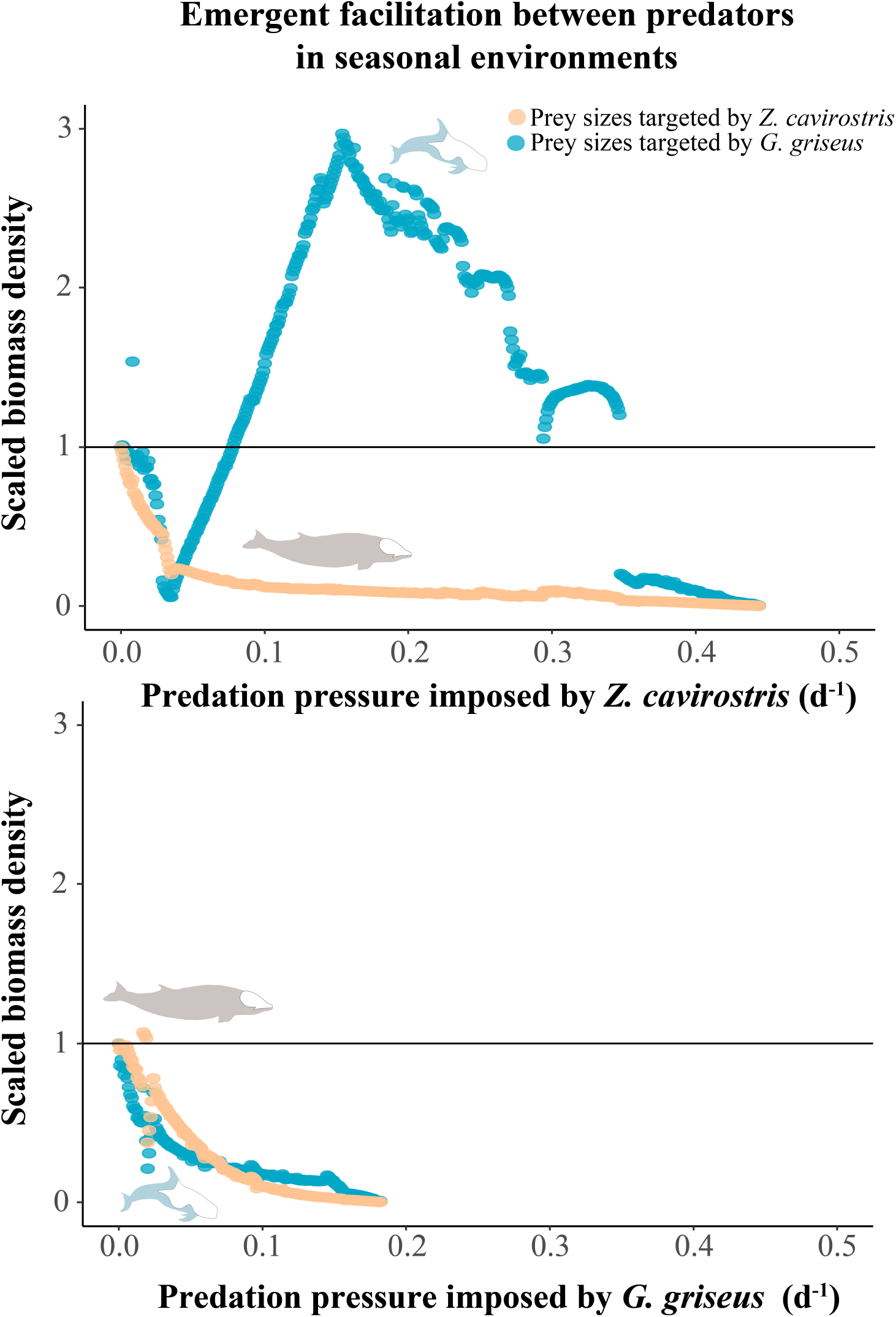
The scope for emergent facilitation by *Z. cavirostris* is larger in seasonal environments. The scaled biomass density of squid prey size classes preferred by *Z. cavirostris* (orange-pink line) and *G. griseus* (blue line) are presented as a function of the predation pressure by each predator (a, b). The scaling is relative to the average prey biomass density when there is no predation by either predator (horizontal line at 1.0 on the y-axis). Emergent facilitation occurs when prey-biomass density exceeds 1.0 for a predator as a function of increased predation pressure by another predator. Seasonality in the model is set to *ψ* = 0.7. Parameters *R*_1max_, *R*_*2*max_, *R*_3max_ were set at 1.5, 1.5, and 2.0 g/L respectively. Other parameters as in Table 1.

## Discussion

Hyperabundant semelparous deep-sea squid play a key trophic role in oceanic ecosystems. Here, we present the first mechanistic exploration of how their populations are regulated. Life-stage specific differences in intraspecific resource competition play a critical role. High levels of intraspecific resource competition experienced as paralarvae in surface waters, and the release from competition following ontogenetic vertical migration as juveniles, drive a pattern of faster growth with increasing size. This mechanism causes periodicity in the patterns of maturation and population dynamics. We did not find evidence for the hypothesis that reduced metabolic costs following ontogenetic migration drive faster growth with increasing size. The faster growth with increasing size resulted in exponential-like individual growth curves, and these can thus be explained as emergent outcomes from ecological (e.g. size-specific resource competition), rather than physiological feedbacks in our model. The resource feedbacks we describe lead to the formation of single-cohort cycles in the squid population with periodical alterations between the presence of paralarvae and small, or maturing juveniles. The structure of the population during these cycles changes depending on the degree of seasonality in zooplankton productivity. Moreover, we find that the population dynamics of *H. reversa* are strongly influenced by size-selective predation at different depths. This size-selective predation can lead to emergent facilitation between the predators *Z. cavirostris* and *G. griseus*, because of their complementary size-specific prey preferences.

These findings identify ecological feedbacks that can drive the unusual, yet poorly understood life history of deep-sea squid. The single cohort cycles we describe for *H. reversa* have been described for other marine and freshwater taxa [45, 46], but represent a new and previously unknown form of population regulation for deep-sea squid. Single-cohort cycles are typically found in species that share a single resource throughout their life cycle and whose foraging efficiency decreases with body size [51]. Classic examples include zooplanktivorous fish [52, 53, 54, 55] and phytoplanktivorous *Daphnia* [56, 57, 58, 59]. In these species, new cohorts entering the population gradually outcompete older cohorts by foraging more efficiently on the shared resource, until a single dominant cohort remains. The life history of *H. reversa* differs from this scenario because individuals switch resources as they develop from paralarvae to juveniles, and again as they move to increasing depths. Thus, our results are not driven by an increase in foraging efficiency with increasing body size. Ontogenetic shifts in diet and habitat are a common strategy observed in many groups of animals to limit intraspecific competition between life stages [60, 61, 62, 63, 64, 65, 66]. Ontogenetic migration close to the seafloor also limits predation risk in squid in the reproductive phase [67]. However, ontogenetic shifts in diet and habitat usually do not lead to single-cohort cycles. These cycles seem to emerge in *H. reversa* because reproductive inflow is so high that some cohorts gradually start to outcompete themselves (by suppressing their own resources below critical levels), until, over time, one cohort remains. In a sense, *H. reversa* populations fall victim to their own reproductive success. This raises the question of why squid populations might maintain such high reproductive investments, if it is followed by intense competition and starvation of offspring. The answer to this question may be found in evolutionary life history theory which [68, 69, 70] predicts that evolution maximizes individual, not group, fitness [70]. Thus, lower reproductive investments would only serve to reduce the fitness of the reproducing individual. Furthermore, investing in producing a large number of small paralarvae, rather than fewer larger ones, is beneficial in semelparous species when offspring survival depends on favourable, but variable conditions over time [71, 72]. For example, in areas where there is strong seasonality in primary productivity, such as in oceanic fronts or upwelling areas [71, 73, 74]. Finally, a recent study suggests that high predation pressure by deep-diving whales, from which deep-sea squid cannot escape by means of physical protection, agility, or schooling, could be a key driver of the fast semelparous life histories of deep-sea squid with high reproductive investment [75].

Deep-sea squid populations are estimated to be highly abundant [76, 77], but a large open question is how we might expect their populations to respond to climate change. We showed that population dynamics of semelparous deep-sea squid are strongly linked to the development of early life stages in surface waters. Global sea surface water temperatures are increasing under climate change [78] and experiments indicate this will increase the growth rate of hatchling cephalopods [79, 80]. These increased growth rates are driven by increased individual food-intake at higher water temperature [81]. *In-situ*, we might thus expect increases in surface water temperatures to result in increased resource demands (and competition) of deep-sea squid paralarvae. However, a big unknown is how changes in sea surface water temperatures will affect the phenology and productivity of their zooplankton prey base [82, 83]. Our model indicates that where zooplankton resource productivity is reduced, we might expect the total biomass of deep-sea squid populations to decrease (Supplemental Figure 4.3). However, in areas where zooplankton productivity will increase, we can expect the total biomass of deep-sea squid populations to increase (Supplemental Figure 4.3). This latter pattern has already been observed in the in the Sea of Japan [84]. Here, rising sea surface water temperatures were correlated to higher zooplankton biomass production and increased commercial squid catches [84]. A second important effect will be the range shifts in zooplankton production under climate change, such as polewards extensions [85], from which deep-sea squid populations might profit in some areas. However, reviews show these range shifts have proven inconsistent and species-specific in zooplankton [83, 86]. It is therefore difficult to predict the expected climate change effects on the future distribution of semelparous deep-sea squid populations.

The responses of deep-sea squid populations to size-selective predation offer fundamental insights into the intricate dynamics shaping species communities in the pelagic deep sea. Research has consistently shown that diverse communities of top predators are key to maintaining ecosystem functioning, productivity, and diversity across trophic levels [87, 88]. However, we also show that the benefits of increased prey availability by size-selective predation are much greater for predators targeting smaller juvenile squid at shallower depths (*G. griseus*) than for those targeting mature individuals in the deeper zone (*Z. cavirostris*). This unexpected outcome could be one of the factors contributing to the fact that *Z. cavirostris* typically holds smaller population sizes (100s), opposed to larger population sizes for *G. griseus* (1000s), specifically in areas where they co-occur [89, 90, 91]. More generally, it suggests that foraging on smaller squid at shallower depths might be a more rewarding and stable strategy for deep-diving cetaceans than specializing on foraging at extreme depths. Still, the diversity of cetacean species that target deeper layers suggests that foraging at these depths remains advantageous. This may be because apart from *H. reversa* there are many other important deep-sea squid prey species in these deeper layers, which we know are consumed by deep divers but that are not accounted for in our model.

We showed that the scope for emergent facilitation strongly increases under seasonal resource dynamics (Figure 6). This outcome indicates we can expect higher population levels, or more species of deep-diving cetaceans co-occurring in seasonal environments. This aligns with the observation that in hotspot areas such as the Azores there are seasonal patterns in the arrival of additional deep-diving cetacean predators with potentially overlapping niches in depth and diet compared to more ‘resident’ species (e.g. *Hyperodon ampullatus*, male *P. macrocephalus*) [90]. We conclude that a combined approach of theoretical, evolutionary, and observation-based studies exploring a fuller predator context are needed to understand how deep-sea ecosystems can support and maintain such diverse communities of top predators. These studies are essential to define a more general theory of the functioning of multi-trophic interactions in pelagic deep-sea ecosystems. We urgently need such theory to better understand and predict how deep-sea communities across trophic levels, including deep-diving cetaceans as the largest top-predators in these systems, will respond in the face of increasing anthropogenic pressures in this era of global change [92, 93, 94, 95].

