## Supplementary Information for "Population regulation in semelparous deep-sea squid is driven by ecological conditions in surface waters and whale predation at depth"

### S1. Ontogenetic depth migration

A visualization of the function describing ontogenetic depth migration.

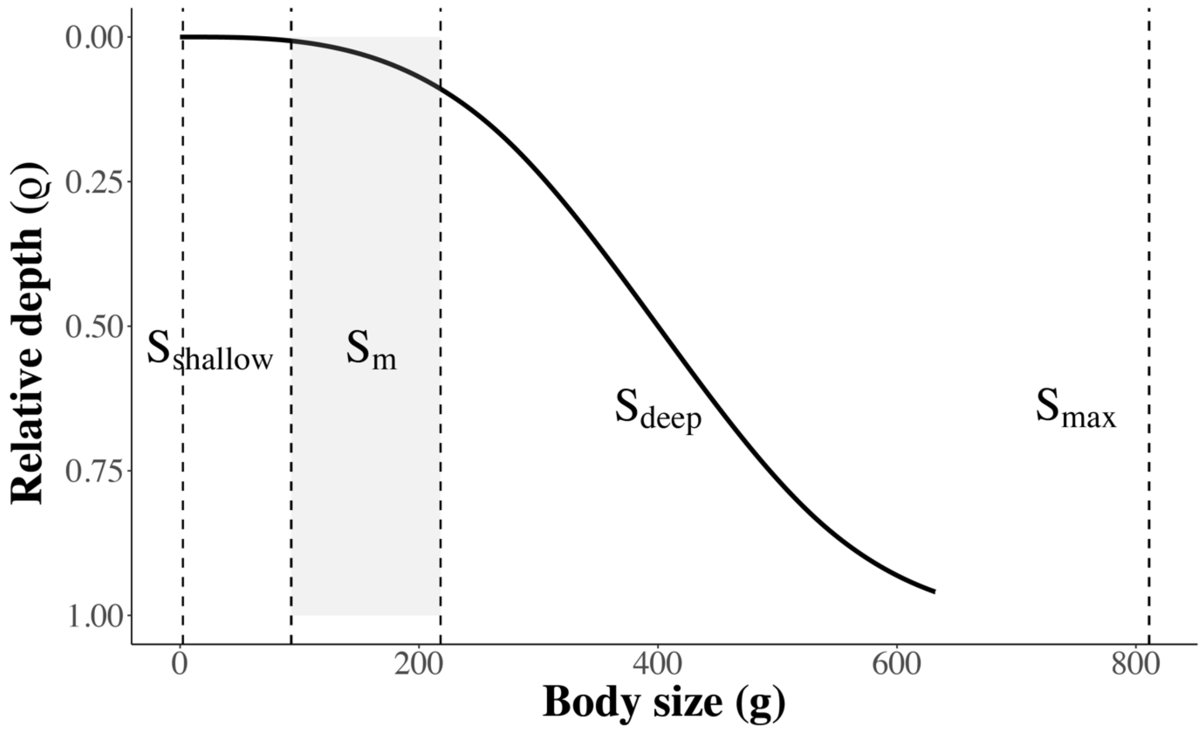


**Supplementary Figure 1.** The depth inhabited by individual *H. reversa* as a function of body size. Where $S_{shallow}$ is the size at which individuals start depth migration, $S_{deep}$ is the size at half the maximum depth migration, and $S_{max}$ is the theoretical maximum body size. The shaded area represents the observed range of maturation sizes *in-situ*, $S_m$. Default parameters values are given in Table 1.

#### S2. Impact of metabolic maintenance

We find no differences in the population dynamics, growth curves, or mass specific growth in *H. reversa* when mass-specific metabolic costs decrease with depth, or when metabolic costs are constant over depth (Supplementary Figure 2).

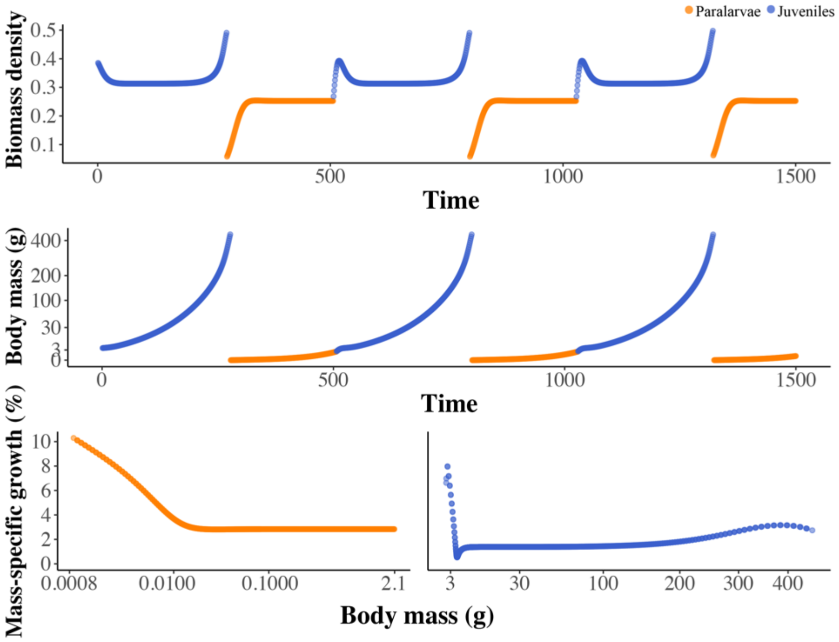


**Supplementary figure 2**. Patterns in time series dynamics (top), growth curves (middle), and mass-specific growth (bottom) when there is no reduction in mass-specific metabolic maintenance costs with depth. Orange colours refers to individuals in the paralarval life stage and blue to individuals in the juvenile life stage. Default parameters values are given in Table 1.

#### S3. Robustness of growth curves

We find the pattern of exponential like growth and underlying increased mass-specific growth with increasing size to be a robust outcome in *H. reversa*. Only a combination of high paralarval mortality ($\mu_j$), low productivity of the secondary nekton resource ($R_3$) at the benthic boundary layer, and an earlier switch to this resource ($S_{deep}$), results in a qualitative reversion where mass-specific growth decreases with increasing size and there is no clear exponential pattern in growth (though growth still does not reach an upper plateau).

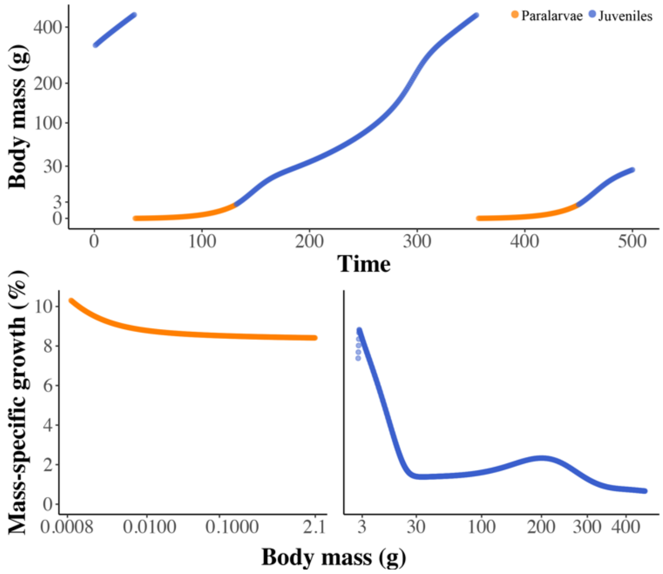


**Supplementary Figure 3.1** Extreme ecological conditions (low R3 resource productivity, high paralarvae mortality, direct switch to benthic resource feeding as juveniles) resulting in a qualitative reversion where growth trajectories become more linear (a) and mass-specific growth decreases with increasing size (b,c). Orange colours refers to individuals in the paralarval life stage and blue to individuals in the juvenile life stage. $R_{3max}$ equals 0.2 gram/L, paralarvae mortality $\mu_j$ equals 0.08 per day, and the size at the switch to the benthic resource equals 200 gram. All other parameters have default values as given in Table 1.

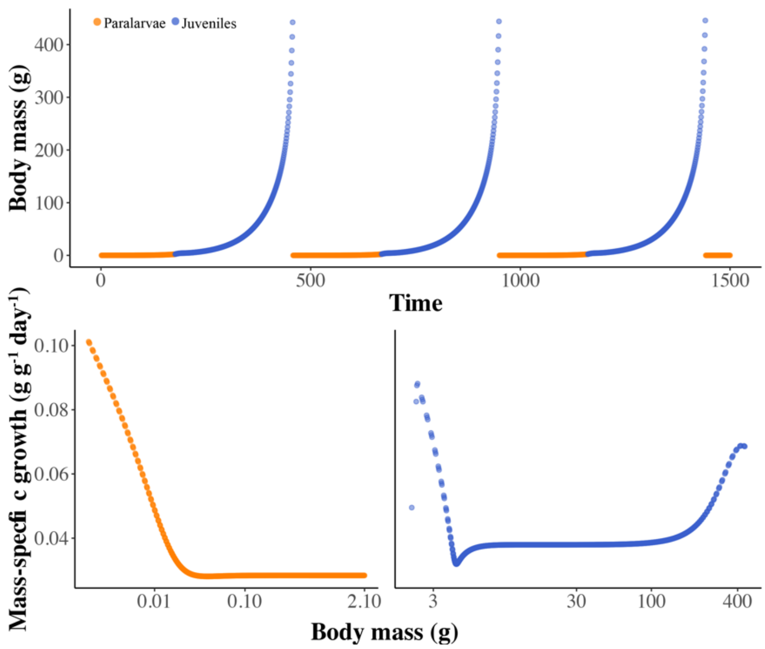


**Supplementary Figure 3.2.** Individual growth curve (a) and mass-specific growth (b, c) for default parameters (Table 1). The same patterns in growth remain as in figure 2 of the main text, but we did not apply a square-root scaling of the y-axis of (a). Because growth spans over seven orders of magnitude, no insight can be gained in the growth in panel (a) during the paralarval stage using an untransformed axis. Orange colours refers to individuals in the paralarval life stage and blue to individuals in the juvenile life stage

### S4. Bifurcation analysis of life history and resource parameters

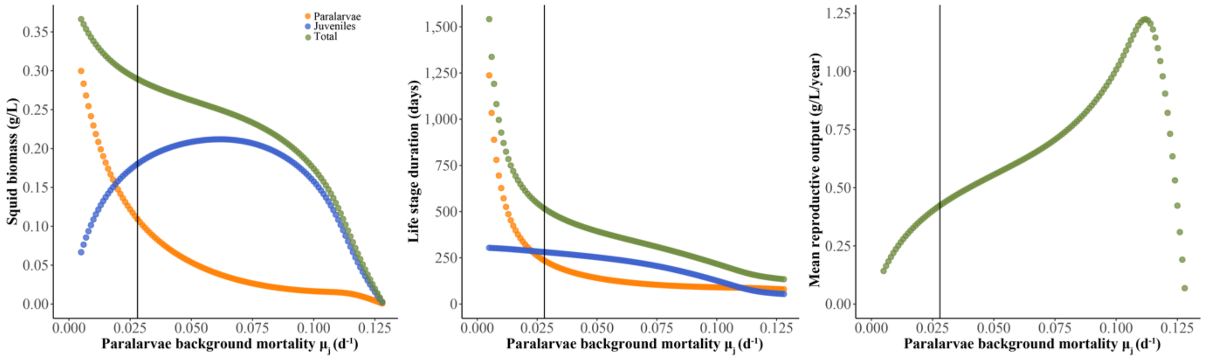


**Supplementary Figure 4.1** Average biomass of the squid life stages (panel a), duration of each life stage (b), and reproductive output (c) as a function of the background mortality rate of paralarvae $\mu_{\mathrm{j}}$. Vertical lines indicate the default background paralarvae mortality rate. Apart from changing paralarval mortality, all other parameters were kept at their default values (Table 1). Orange colours refers to individuals in the paralarval life stage, blue to individuals in the juvenile life stage, and green shows the total of the two life stages.

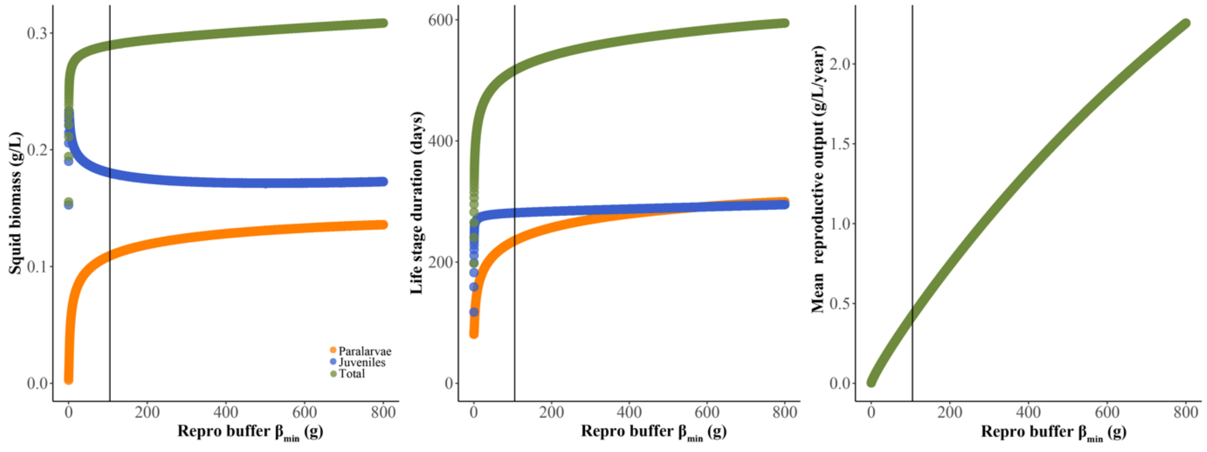


**Supplementary Figure 4.2** Average biomass of the squid life stages (a), duration of each life stage (b), and reproductive output (c) as a function of changing reproductive buffer mass $\beta_{\mathrm{min}}$ in grams. Apart from changing parameter $\beta_{\mathrm{min}}$, all other parameters were kept at their default values (Table 1). Orange reflects biomass of individuals in the paralarval life stage, blue reflects biomass of individuals in the juvenile life stage, and green reflects the total of the two life stages.

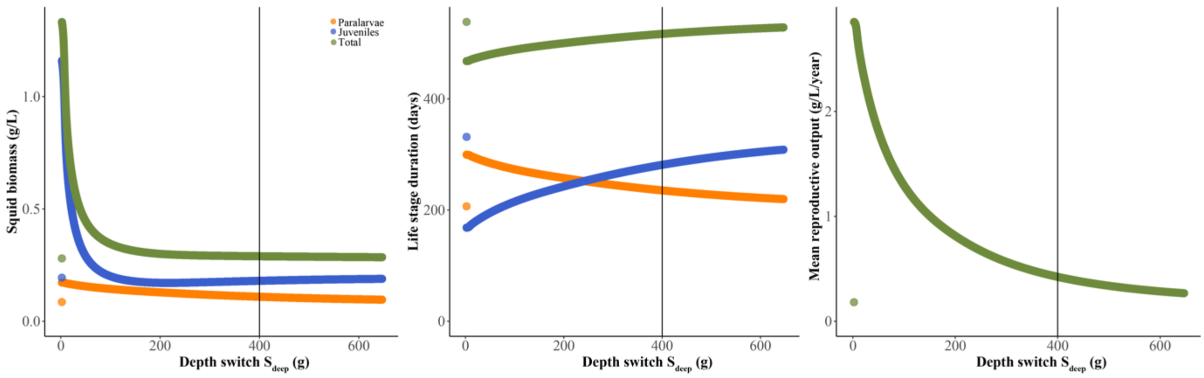


**Supplementary Figure 4.3** Average biomass of the squid life stages (panel a), duration of each life stage (b), and reproductive output (c) as a function of changing the size at half the maximum depth migration ($S_{deep}$). Orange colours refers to individuals in the paralarval life stage, blue to individuals in the juvenile life stage, and green the total of the two life stages. Apart from changing parameter $S_{deep}$ , all other parameters were kept at their default values (Table 1).

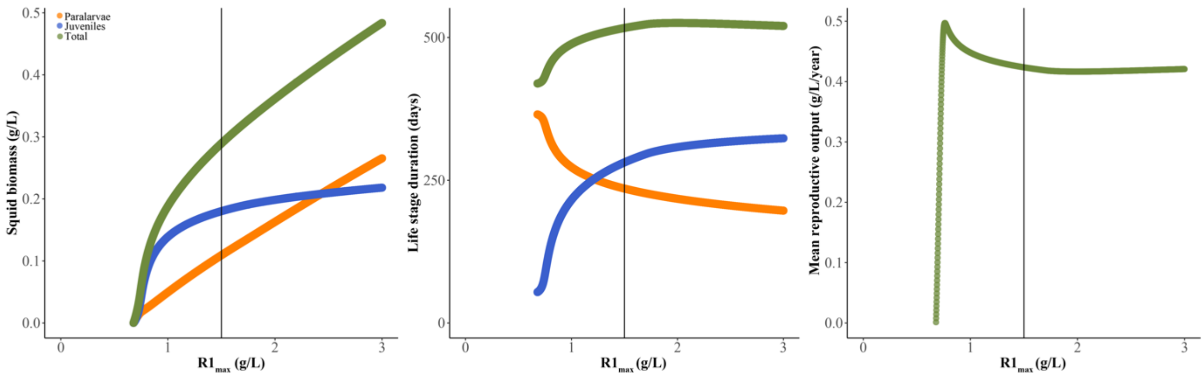


**Supplementary Figure 4.3** Average biomass of the squid life stages (a), duration of each life stage (b), and reproductive output (c) as a function of changing zooplankton resource productivity ($R_{1max}$). Orange colours refers to individuals in the paralarval life stage, blue to individuals in the juvenile life stage, and green the total of the two life stages. Apart from changing parameter $R_{1max}$, all other parameters were kept at their default values (Table 1).

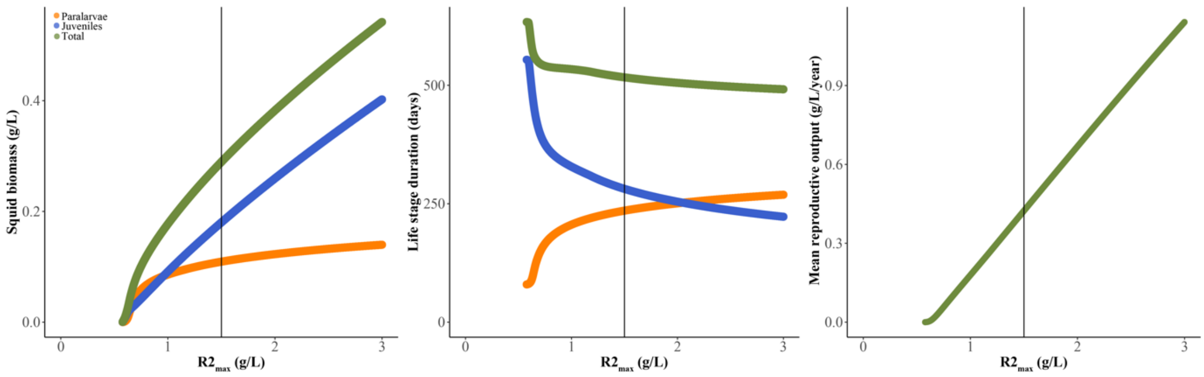


**Supplementary Figure 4.5** Average biomass of the squid life stages (a), duration of each life stage (b), and reproductive output (c) as a function of changing the first nekton resource productivity ($R_{2\max}$). Orange colours refers to individuals in the paralarval life stage, blue to individuals in the juvenile life stage, and green the total of the two life stages. Apart from changing parameter $R_{2\max}$, all other parameters were kept at their default values (Table 1).

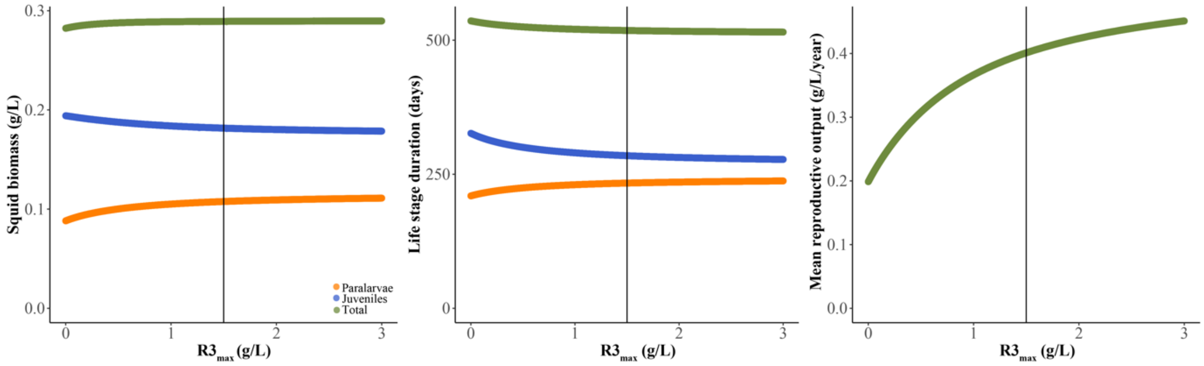


**Supplementary Figure 4.6** Average biomass of the squid life stages (a), duration of each life stage (b), and reproductive output (c) as a function of the productivity of the second nekton resource ($R_{3\max}$). Orange colours refers to individuals in the paralarval life stage, blue to individuals in the juvenile life stage, and green the total of the two life stages. Apart from changing parameter $R_{3\max}$, all other parameters were kept at their default values (Table 1).

### S5. Seasonal versus aseasonal population dynamics

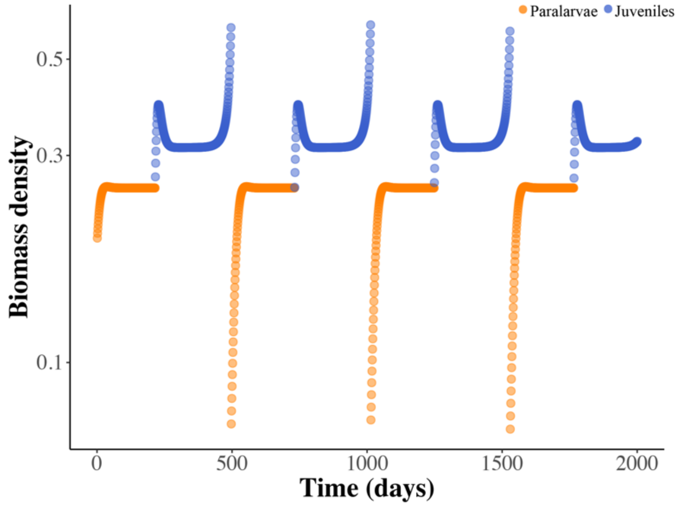


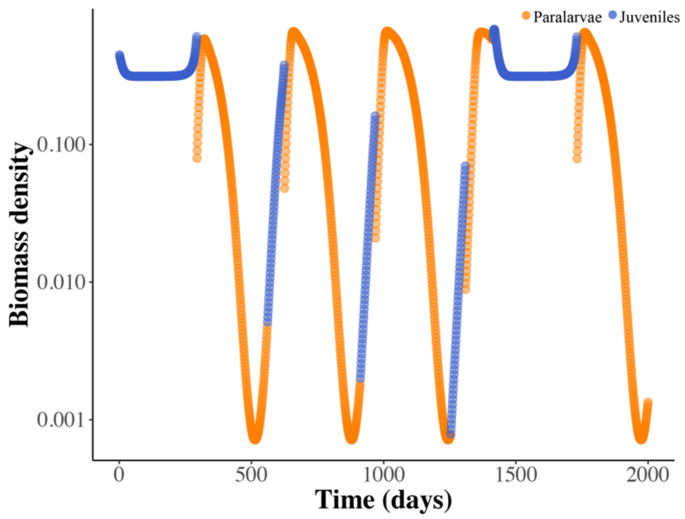


**Supplementary Figure 5.** Population dynamics of *H. reversa* as a function of time with no seasonality in zooplankton productivity ($\varphi$ = 0; top), and high seasonality in zooplankton resource productivity ($\varphi$ = 0.8; bottom). The average population biomass is highest in the juvenile stage (blue line) in aseasonal environments and highest in the paralarval life stage (orange line) in seasonal environments. In both cases population dynamics remain characterized by single cohort cycles, but in seasonal environments the bottleneck in the paralarval life stage is lifted. All other parameters were kept at their default values (Table 1).

#### S6. Size selective predation and squid time series dynamics (aseasonal)

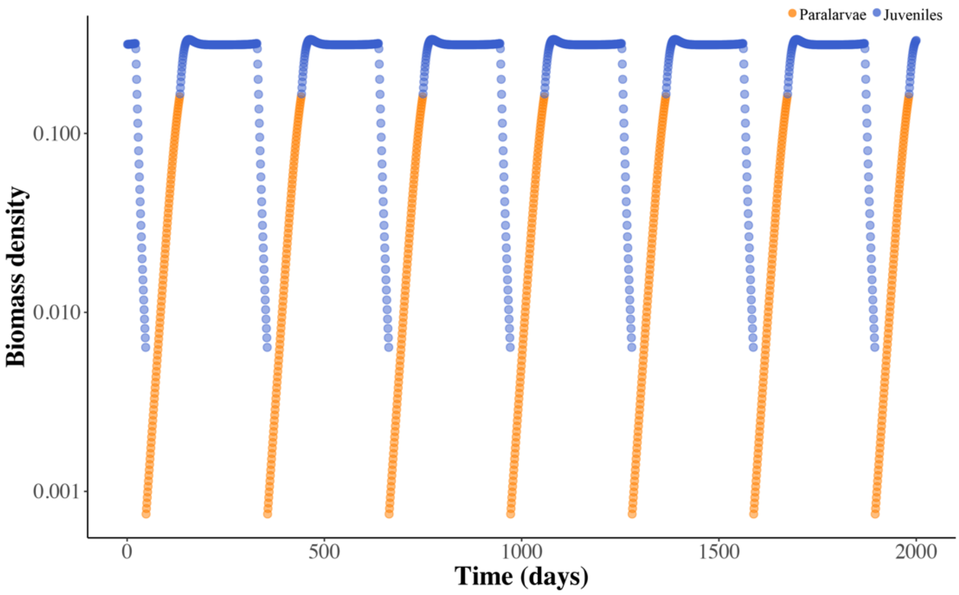


**Supplementary Figure 6.1**. Time series population dynamics of squid density with additional size-selective mortality imposed by Cuvier's beaked whale. The size selective mortality rate ($\mu_c$) is set at 0.2 per day, all other parameters as in Table 1. Orange line depicts biomass density in the paralarval life stage, and the blue line the biomass density in the juvenile life stage.

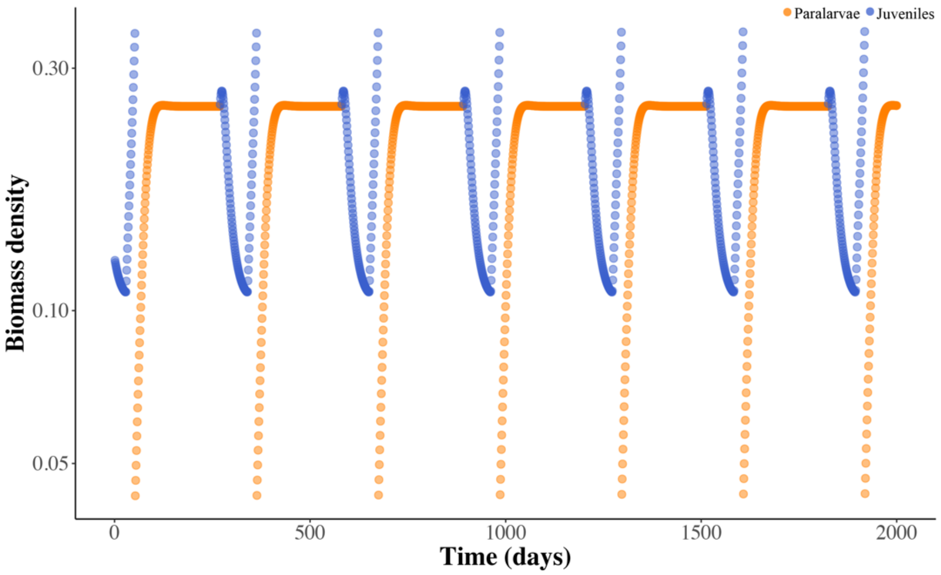


**Supplementary Figure 6.2**. Time series population dynamics of squid density with additional size-selective mortality imposed by Risso's dolphin. The size selective mortality rate ($\mu_c$) is set at 0.2 per day, all other parameters as in Table 1. Orange line depicts biomass density in the paralarval life stage, and the blue line the biomass density in the juvenile life stage.

#### S7. Size selective predation and squid time series dynamics (seasonal)

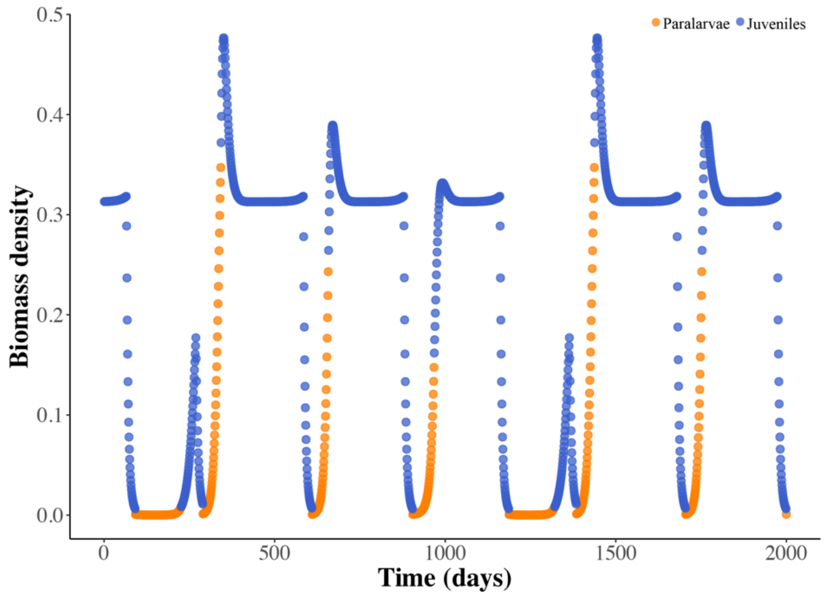


**Supplementary Figure 7.1**. Time series population dynamics of squid density with additional size-selective mortality imposed by Cuvier's beaked whale in seasonal environments. The size selective mortality rate ($\mu_c$) is set at 0.2 per day and seasonality in zooplankton productivity ($\varphi$) at 0.8, all other parameters as in Table 1. Orange line depicts biomass density in the paralarval life stage, and the blue line the biomass density in the juvenile life stage.

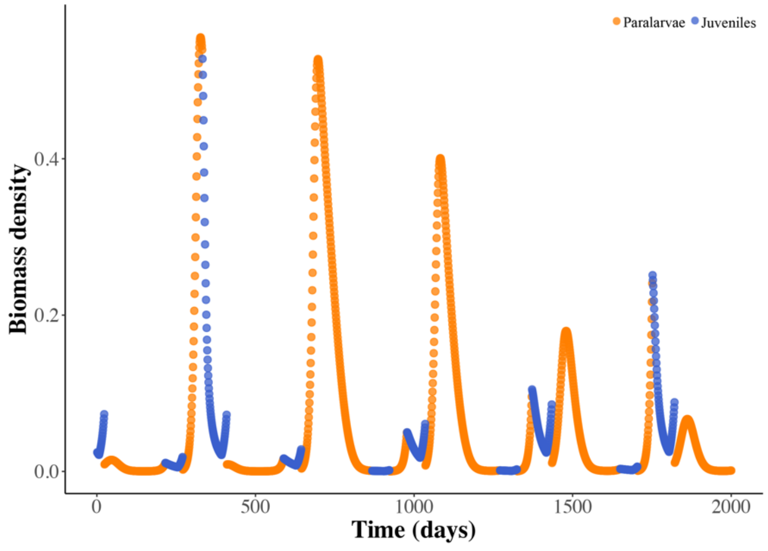


**Supplementary Figure 7.2**. Time series population dynamics of squid density with additional size-selective mortality imposed by Risso's dolphin in seasonal environments. The size selective mortality rate ($\mu_g$) is set at 0.1 per day and seasonality in zooplankton productivity ($\varphi$) at 0.8, all other parameters as in Table 1. Orange line depicts biomass density in the paralarval life stage, and the blue line the biomass density in the juvenile life stage.
